# Evolution under vancomycin selection drives divergent collateral sensitivity patterns in *Staphylococcus aureus*

**DOI:** 10.1101/2023.11.30.569373

**Authors:** Kyle J. Card, Dena Crozier, Arda Durmaz, Jason Gray, Justin Creary, Amira Stocks, Jeff Maltas, Robert A. Bonomo, Zachary D. C. Burke, Jacob G. Scott

## Abstract

*Staphylococcus aureus* bacteremia is typically treated empirically with vancomycin, with therapy later tailored based on susceptibility results. However, these tests occur before vancomycin exposure and do not account for adaptation during empiric treatment that can alter *S. aureus*’ susceptibility to first-line drugs. To investigate these collateral drug responses, we experimentally evolved 18 methicillin-susceptible *S. aureus* (MSSA) populations under increasing vancomycin concentrations until they achieved intermediate resistance. Genomic sequencing revealed two distinct adaptive pathways characterized by mutations in the WalKR regulon, affecting cell wall metabolism, or *rpsU*, impacting translational stress responses. These pathways correlated with divergent collateral sensitivity profiles to first-line antibiotics. By developing a Collateral Response Score (CRS), we quantified the probability and magnitude of these responses, demonstrating that evolutionary dynamics critically influence resistance outcomes. Our findings suggest a probabilistic approach to antimicrobial therapy, advocating for rapid genomic diagnostics alongside susceptibility testing to better anticipate and respond to evolutionary changes.

**Significance:** Antibiotic treatment can influence bacterial evolution, altering the effectiveness of subsequent therapies by inducing collateral resistance or sensitivity. This study reveals that evolution toward vancomycin-intermediate resistance in the pathogen *Staphylococcus aureus* proceeds through at least two distinct evolutionary pathways: one characterized by alterations in cell wall metabolism and another by changes in global stress response. These adaptive trajectories result in contrasting collateral sensitivities to first-line antibiotics. By introducing the Collateral Response Score, we assess the uncertainty in these outcomes, providing a probabilistic framework to evaluate how past antibiotic exposure shapes future treatment responses. Further validation studies are needed; however, we believe that improved forecasting of pathogen evolution can enhance antibiotic stewardship, inform therapeutic decisions, and ultimately improve patient outcomes.

## Introduction

*Staphylococcus aureus* is a Gram-positive pathogen often responsible for severe infections, including endocarditis, osteomyelitis, soft tissue infections, and device-associated infections (1–4). It typically enters the bloodstream through various routes, often via a wound, surgical site, or catheter, frequently leading to bacteremia (1, 3). When clinicians suspect a severe Staphylococcal infection, they commonly initiate treatment with broad-spectrum antibiotics, such as vancomycin, to address the possibility of methicillin-resistant *S. aureus* (MRSA). Blood culture results can take several days to return, at which time therapy may be adjusted based on the detected organism and its susceptibility profile. Currently, antistaphylococcal penicillins, including nafcillin and oxacillin, are recognized treatments for methicillin-susceptible *S. aureus* (MSSA) infections, and clinicians may administer these drugs if cultures exclude MRSA. However, we ask whether favorable susceptibility results alone provide a complete picture. This question rests on two key considerations.

First, the evolution of vancomycin-intermediate resistance during empiric treatment may affect *S. aureus*’ susceptibility to other drugs (i.e., collateral drug responses). Intermediate resistance evolves via mutations in a diverse set of genes (5–8) and is associated with persistent infections and reduced treatment success (8). These mutations may alter MSSA’s sensitivity to subsequent first-line therapies, even in the absence of prior exposure. For example, a large cohort study showed that the in-hospital mortality rate of cloxacillin-treated patients with MSSA correlated with vancomycin minimum inhibitory concentration (MIC) (9). MSSA should be susceptible to both cloxacillin and methicillin because these drugs have the same cellular target. Prior vancomycin adaptation is one possible contributing factor to these poor clinical outcomes.

Second, antibiotic susceptibility test results reflect a single snapshot in time, often before empiric antibiotic therapy, and are thus a lagging indicator of phenotypic state and agnostic to evolution. In current practice, clinicians sample a patient’s blood and then administer broad-spectrum antibiotics. They isolate the causative agent of the infection from the blood cultures and rely on susceptibility tests to inform subsequent therapy. However, these tests provide information about the pathogen’s susceptibilities before treatment and cannot account for any potential changes in its antibiotic response arising from intervening evolution under empiric therapy. Moreover, susceptibility results only reflect a subset of the bacterial population existing in a patient. Initial culture results may indicate that the infection is susceptible to an antistaphylococcal penicillin; however, this recommendation may be inaccurate, given that the infection has had the opportunity to evolve for several days (several hundred bacterial generations) in this new environment, potentially altering antibiotic susceptibility profiles in various ways (10, 11).

To address these issues, we investigated how empiric therapy impacts collateral drug sensitivities by experimentally evolving replicate MSSA populations under increasing vancomycin concentrations until they reached intermediate resistance levels. The populations exhibited diverse collateral responses to several first-line antibiotics. However, the lines broadly followed two adaptive routes under vancomycin selection, rendering some drug tradeoffs explicable by genetic background. Ideally, a treatment regimen for MSSA would include drugs that the infection has a high chance of being susceptible to, given its prior vancomycin exposure. Thus, background-specific interactions complicate therapy yet underscore the need for rapid and accurate genomic sequencing, alongside standard antimicrobial susceptibility tests, to inform therapeutic decisions. In that spirit, we propose the Collateral Response Score (CRS), a standardized metric that provides information about the direction and magnitude of changes in MIC.

Taken together, our results underscore the uncertainty and risk of not accounting for evolution when making therapeutic decisions. Instead, clinicians should anticipate that infections will evolve under empiric therapy, which might affect their susceptibility to first-line drug treatment. Our study, therefore, highlights the complexities of bacterial evolution and emphasizes that we should consider susceptibilities in a probabilistic light, just as we do with other inherently stochastic systems.

## Results and Discussion

### Evolution of vancomycin-intermediate resistance in MSSA

To model Staphylococcal evolution under empiric vancomycin therapy, we established 18 replicate MSSA populations from an ancestral clone of *S. aureus* subsp. *aureus* Rosenbach, a clinical isolate that is an international quality control standard. We transferred each population daily into a growth medium containing vancomycin. We gradually increased the concentration until the lines reached intermediate resistance levels (**Fig. 1*A***). Additionally, 87 control lines were established from the same ancestral strain and propagated under identical conditions except in the absence of vancomycin. It is important to note that controls in experimental evolution differ from those used in other fields of biology. When the null hypothesis is that an expected effect will or will not occur based on the presence of an experimental perturbation, single or paired controls make sense. Here, however, our null hypothesis is not that the control lines will have a specific, single outcome different from the experimental conditions, but rather that they will display an ensemble of outcomes that differs from those of the experimental group. In this study, we therefore examined the differences in the distributions of these mutational outcomes.

**Fig. 1.**
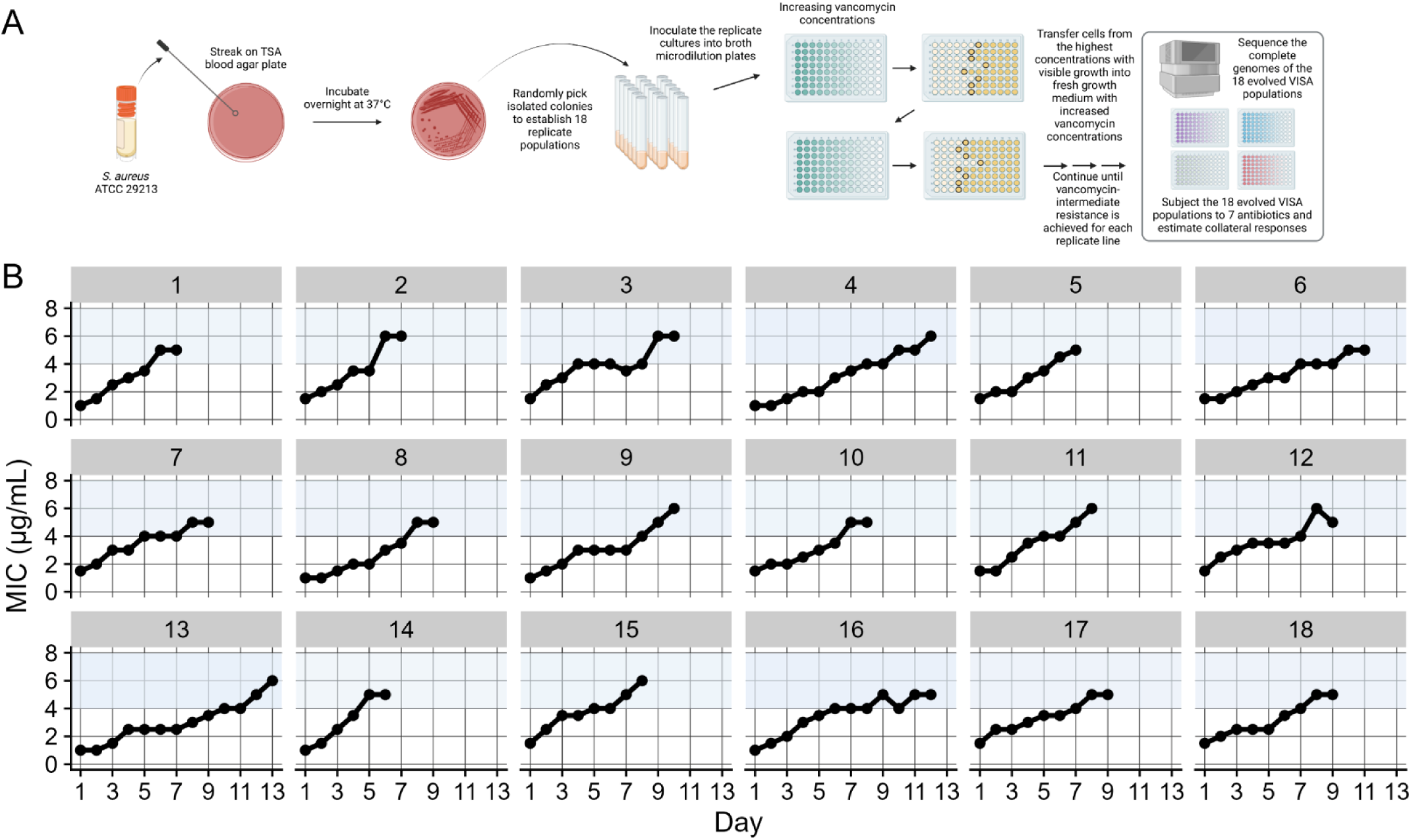
Schematic illustration of our study design and evolution of intermediate vancomycin resistance. (*A*) We established 18 independent populations from a methicillin-susceptible *S. aureus* (MSSA) strain and experimentally evolved them under increasing vancomycin concentrations until they reached intermediate resistance levels. Additionally, we established 87 replicate control lines from the same ancestral strain and transferred them under identical conditions except in the absence of vancomycin (not shown). We then performed whole-genome sequencing on all evolved vancomycin-adapted and control populations, and examined the susceptibilities of the vancomycin-adapted lines to seven drugs used in the treatment of MSSA. **(*B*)** By day 13 of the evolution experiment, all vancomycin-adapted populations exhibited intermediate resistance, as defined by the Clinical and Laboratory Standards Institute (CLSI), characterized by a minimum inhibitory concentration (MIC) between 4 and 8 µg/mL (blue-shaded regions).

The 18 vancomycin-exposed lines had an initial average MIC of 1.3 µg/mL (standard deviation, *SD* = 0.3) and subsequently evolved to an average MIC of 5.4 µg/mL (*SD* = 0.5) (**Fig. 1*B***). The time required for the lines to increase their MIC varied significantly. It took an average of 9 days (*SD* = 2), with population 14 achieving vancomycin-intermediate resistance in just six days, while population 13 required 13 days to reach the same resistance level.

### Populations follow distinct adaptive pathways under vancomycin selection

Next, we sequenced the ancestor and experimentally evolved populations to investigate the genetic changes associated with vancomycin-intermediate resistance. The resistant and control lines had 4,921 mutations at a frequency of ≥5% (**Fig. 2*A***): 81.9% were single-base substitutions, and 14% were intergenic mutations located within 150 bp upstream of a gene, suggesting that they affect regulation. Structural variants were rare, and gene amplification was almost non-existent, likely due to the low abundance of repeat regions that mediate tandem amplifications (12).

**Fig. 2.**
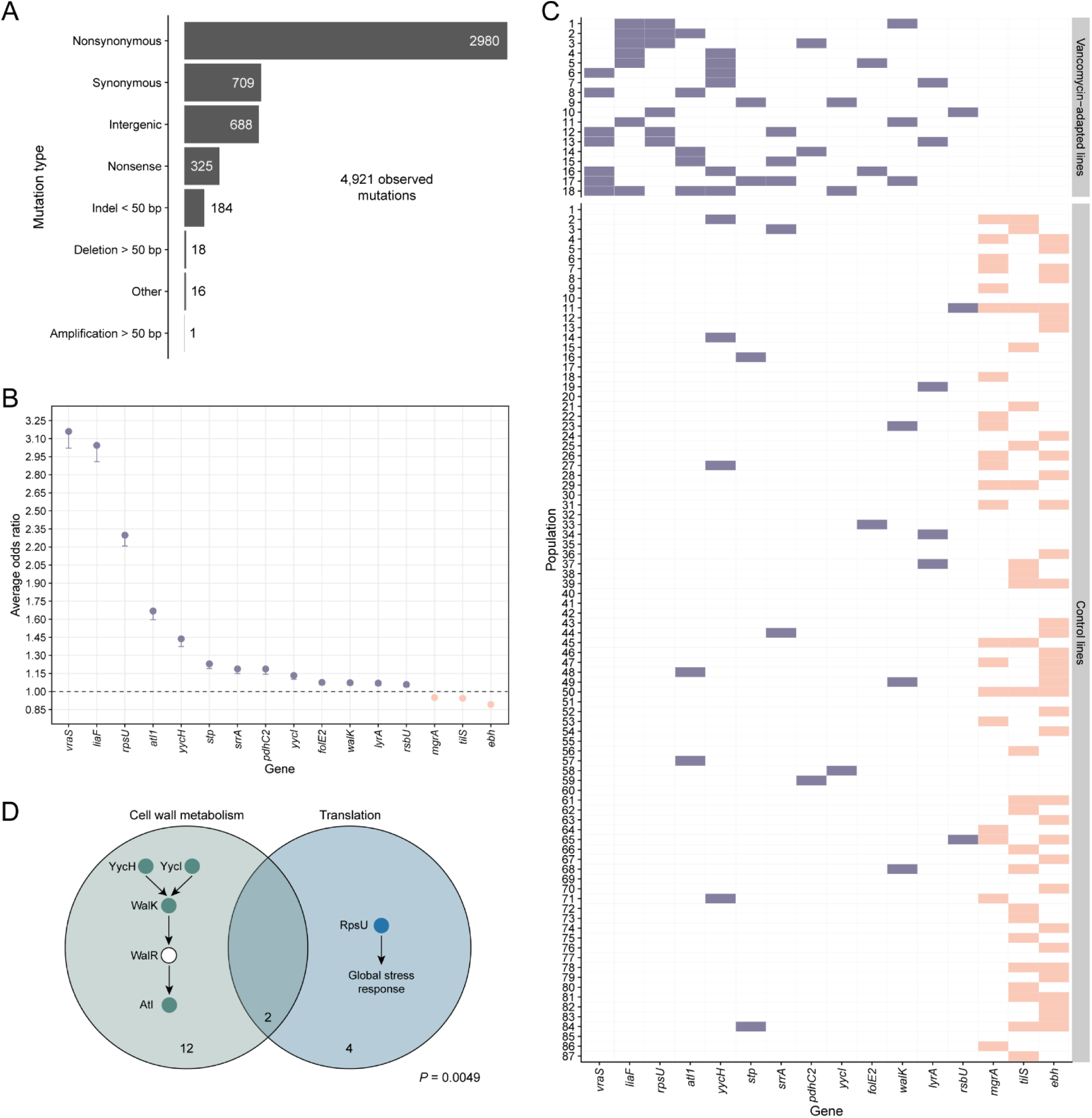
Identification of mutated genes and pathways in experimentally evolved vancomycin-intermediate *S. aureus* (VISA) populations. (*A*) Classification of mutations by type. Numbers indicate the mutation counts for each category. **(*B*)** The average odds ratios for genes significantly associated with vancomycin exposure (violet-filled points) and control conditions (salmon-filled points) were evaluated across 250 iterations of a multivariate logistic regression procedure (*Materials and Methods*). Error bars illustrate 95% confidence intervals, calculated from the model coefficients and subsequently exponentiated. The horizontal dashed line indicates an odds ratio of one. Point estimates are arranged by mean odds ratio. **(*C*)** Identification of genes containing qualifying mutations, arranged by their mean odds ratio and colored in the same manner as panel B. **(*D*)** A Venn diagram illustrating the extent of overlap between the WalKR regulon and *rpsU* mutations in the evolved vancomycin-intermediate *S. aureus* (VISA) populations. Filled circles in each pathway denote mutations impacting those proteins, while the numbers indicate the total number of lines affected.

To assess the impact of vancomycin exposure on genomic parallelism, we compared the gene-level similarity of mutations among independent lineages that evolved under vancomycin selection with those that evolved under permissive, drug-free conditions. As described in the *Materials and Methods*, we computed Dice’s similarity coefficient for each pair of populations using only the 3,812 qualifying mutations that unambiguously affected a single gene. The vancomycin-adapted (VISA) lines exhibited a higher average pairwise similarity (*S_v_* = 0.063) compared to the control lines (*S_c_* = 0.029); that is, on average, two independently evolved populations under vancomycin selection shared 6.3% of their mutated genes, whereas lines that evolved without drug exposure shared only 2.9%. A permutation test confirmed that this difference was significant (*P* < 0.0001), demonstrating that vancomycin selection decisively promotes greater genomic parallelism relative to permissive conditions.

Multivariate logistic regression identified several genes associated with vancomycin exposure. First, *vraS* and *liaF* genes were the most frequently mutated in our lines (**Fig. 2*B***). They occurred in seven out of 18 VISA populations but were absent in all 87 control lines (**Fig. 2*C***). LiaF and VraS regulate the cell membrane stress response (13, 14) and are prevalent in clinical VISA isolates (15). Second, *rpsU* mutations targeting the 30S ribosomal subunit occurred in six VISA lines, and they were also absent in any control population. Ribosomal perturbation can activate stress response in some bacteria. For example, *Listeria monocytogenes rpsU* mutants exhibit upregulated σ^B^-mediated stress genes and slower growth, trading fitness for broad stress resistance (16). In our lines, *rpsU* mutations may similarly impact global regulation, resulting in an elevated cell membrane stress response and reduced cell wall turnover, which can increase vancomycin and daptomycin resistance (12, 17). Indeed, *rpsU* deletion mutants often require compensatory mutations to restore fitness, consistent with this gene’s role in core cellular processes (16). Third, *yycH*, *yycI*, *walK*, and *atl* mutations repeatedly occurred in our evolved VISA lines. These genes comprise the WalKR regulon, an essential cell-wall metabolic pathway in *S. aureus* (**Fig. 2*D***) (18, 19). Loss-of-function mutations in any of these regulon genes cause increased cell wall teichoic acid content, resulting in a thicker and more resistant cell wall against vancomycin and daptomycin (19, 20). This finding is consistent with our results and an earlier study, which showed that mutations in *walKR* cause increased resistance to these antibiotics in clinical VISA strains (5). Together, these results illustrate that vancomycin selection promoted strong genomic parallelism by driving repeated mutations in a limited set of genes primarily involved in cell wall metabolism and general stress response, raising the question of whether evolutionary constraints influenced these adaptive trajectories.

Given these observations, we next investigated whether selection for mutations in specific pathways constrained the evolution of mutations in others, specifically, whether interactions between mutations shaped distinct adaptive trajectories. Bayesian latent class analysis (BLCA) — which considered all vancomycin-associated genes that occurred at least three times across lines and without bias toward predefined functional pathways (e.g., the WalKR regulon) — identified two distinct clusters among the 18 VISA populations, both defined by shared gene-level mutations (**Fig. S2**). Notably, all populations in Cluster 2 evolved mutations in *yycH* and uniformly lacked *rpsU* mutations, whereas six of the twelve populations in Cluster 1 harbored *rpsU* mutations without co-occurring *yycH* alterations. Although the mutual exclusivity between *yycH* and *rpsU* mutations was marginally nonsignificant by Fisher’s exact test (*P* = 0.0537), these findings suggest genomic divergence.

Two distinct adaptive strategies became apparent, however, when we extended our analysis to the functional level, grouping genes by shared biological role. One adaptive strategy modulated cell wall architecture via WalKR regulon mutations, while the other involved a broader reprogramming of translational stress-response pathways mediated by *rpsU* mutations (**Fig. 2*D***). In fact, near complete mutual exclusivity was observed between populations with WalKR regulon mutations and those with *rpsU* mutations: 14 and six lines harbored WalKR regulon and *rpsU* mutations, respectively, with an overlap in only two lines. A Fisher’s exact test confirmed this negative association (*P* = 0.0049). Moreover, 13 of the 14 regulon-mutated lines had a single mutation in this pathway, while the remaining population had three mutations. The probability of this distribution occurring by chance is < 0.0001. WalKR is the only essential two-component system in *S. aureus*, as it controls the expression of enzymes that maintain cell wall architecture during growth and division (18, 20–22). Thus, the partial disruption of this pathway likely imposes a cost on *S. aureus* growth, possibly explaining the rare co-occurrence of WalKR regulon mutations; their combined effect on fitness may be overly deleterious.

In summary, our findings demonstrate that MSSA employed at least two adaptive strategies, balancing the evolution of vancomycin resistance with maintaining core cellular functions. The distinct genomic routes — centered on cell wall remodeling via WalKR regulon mutations and translational reprogramming through *rpsU* alterations — underscore possible sign epistasis that limits the co-occurrence of these mutations. Future work might focus on genomic reconstructions of these evolved mutations into the ancestral background to better examine their epistatic relationships. This divergence is particularly striking, considering that the replicate populations descended from a single ancestor. This result complements previous studies, in which *Escherichia coli* lineages from different genotypes followed divergent paths to increased resistance (23, 24). Finally, our results are consistent with a recent study by Fait and colleagues (25), who evolved 10 independent MRSA populations under vancomycin selection to intermediate resistance and observed notable divergence in mutational profiles; some populations acquired walK mutations, while others evolved mutations in *vraF*.

### Divergent genetic pathways under vancomycin exposure led to varied collateral responses

We then examined how evolution under vancomycin selection affected the susceptibility of VISA populations to commonly used first-line antibiotics. These so-called collateral drug responses are widely observed in bacteria (2, 10, 11, 26–32) and cancer (33–36), and may impact therapeutic outcomes (26). We compared the MICs of the ancestral clone against the 18 evolved populations in cefazolin, clindamycin, daptomycin, gentamicin, meropenem, nafcillin, and trimethoprim-sulfamethoxazole. These antibiotics are suggested treatments for MSSA bacteremia (27) and endocarditis (2, 28). They also have diverse mechanisms of action (**Fig. 3*A***). For each antibiotic, we quantified the collateral response of an evolved population as the difference in its log_2_-transformed MIC relative to the ancestral clone. A population is collaterally resistant when its MIC is higher than the ancestral MIC and collaterally sensitive when it is lower.

**Fig. 3.**
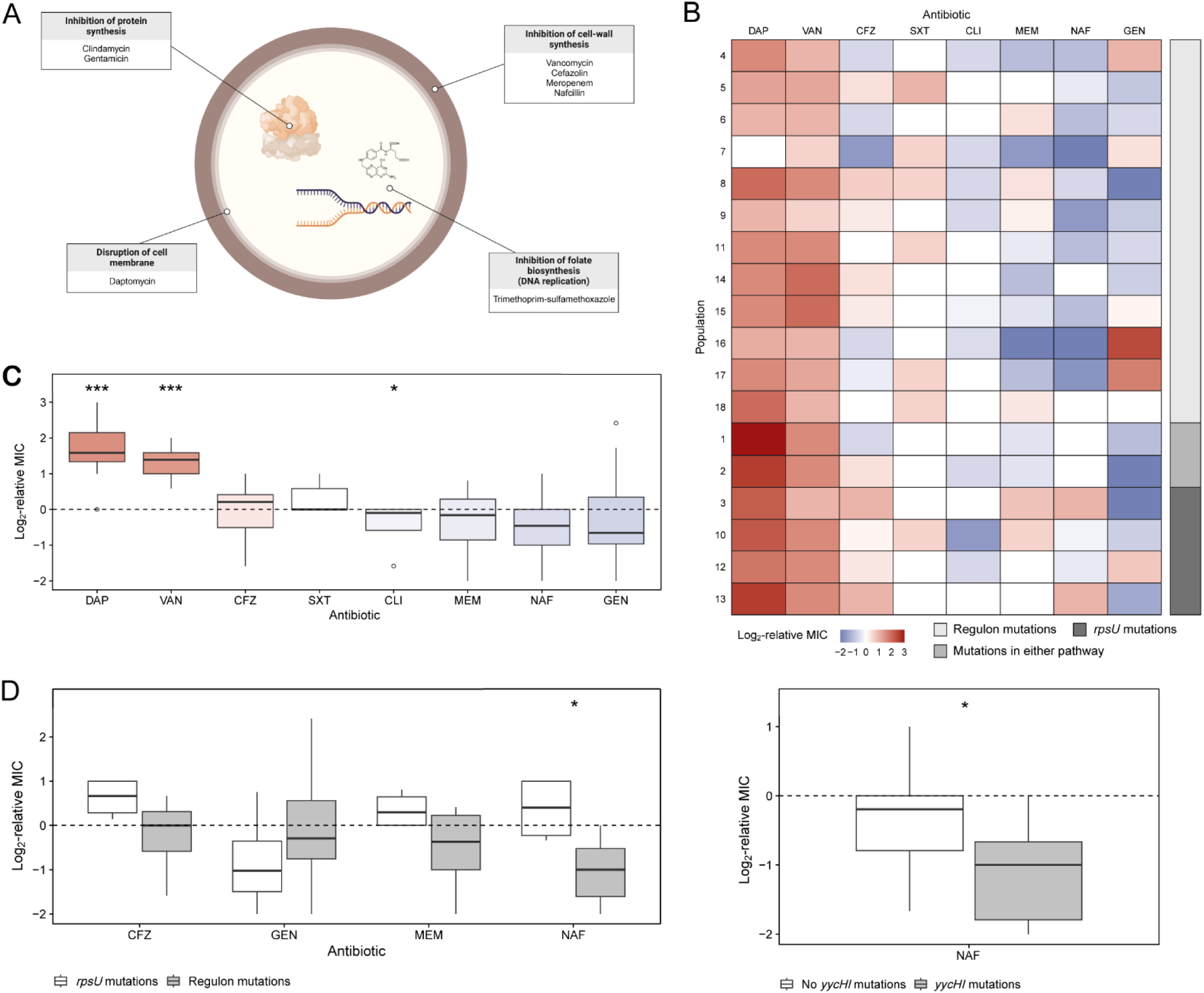
Evolved vancomycin-intermediate *S. aureus* (VISA) lines exhibit diverging collateral sensitivity to cefazolin and nafcillin. (*A*) Bacterial targets and mechanisms of action of each drug. **(*B*)** Collateral resistance (red) or sensitivity (blue) of the 18 evolved VISA lines to each drug expressed as the log_2_-transformed minimum inhibitory concentration (MIC) relative to the ancestor. The lines are ordered by targeted pathway. **(*C*)** Boxplot representing the median and interquartile range of collateral responses in daptomycin (DAP), vancomycin (VAN) (the selecting drug), cefazolin (CFZ), trimethoprim-sulfamethoxazole (SXT), clindamycin (CLI), meropenem (MEM), nafcillin (NAF), and gentamicin (GEN). We ordered the eight antibiotics in decreasing order of median collateral resistance. **(*D*)** Variation in collateral responses between lines with WalKR regulon or *rpsU* mutations and with and without *yycHI* mutations.

The VISA populations exhibited repeated and significant collateral responses to two antibiotics (**Fig. 3*B***). First, 17 out of 18 lines had significantly increased daptomycin resistance relative to the ancestor, as determined by a two-tailed Mann-Whitney test with *P*-values adjusted using the Benjamini-Hochberg method (29) (adjusted *P* = 0.0005). Notably, nine of these lines had MICs above the clinical resistance breakpoint for this phenotype (30) despite no prior exposure to this drug. In contrast, no MIC met or exceeded the clinical resistance threshold to any other drug in our study. Bacteria evolve thicker cell walls under vancomycin selection that hinder daptomycin entry (31–34), likely explaining this finding as well as clinical accounts of collateral resistance (35). Second, the VISA lines were consistently collaterally sensitive to clindamycin (adjusted *P* = 0.0468). Despite these encouraging results, clindamycin monotherapy is associated with higher rates of recurrent infection in MSSA endocarditis because this drug is bacteriostatic (36). Nonetheless, clindamycin is effective when given with trimethoprim-sulfamethoxazole (37).

The evolved populations displayed a range of collateral responses to the five remaining antibiotics, with no significant changes in either direction. We hypothesized that this variation was explained by differences in the genomic profiles that evolved during vancomycin selection. To investigate this possibility, we used point biserial tests to evaluate the associations between collateral response estimates and the presence or absence of mutations. On balance, WalKR regulon mutations were significantly correlated with nafcillin collateral sensitivity (Benjamini-Hochberg adjusted *P* = 0.0178), and *yycHI* mutations alone were sufficient to explain this phenotype (*P* = 0.0372) (**Fig. 3*D***). In contrast, *rpsU* mutations were significantly correlated with cefazolin and nafcillin cross-resistance (adjusted *P* = 0.0379) (**Fig. 3*D***).

Our findings, however, may depend on the specific environmental conditions under which antibiotic susceptibilities were assessed. For instance, Machado and colleagues (38) evolved MRSA populations under different media conditions — cation-adjusted Mueller-Hinton broth (CA-MHB) and Roswell Park Memorial Institute (RPMI) medium — to simulate bacteriological and physiological conditions, respectively. Their results indicated trade-offs between environmental conditions, with lines evolved in CA-MHB showing increased vancomycin resistance in RPMI, but not vice versa. This result suggests that the susceptibility profiles observed in our VISA populations, measured in CA-MHB, may also differ under physiologically relevant conditions. Additionally, Machado and colleagues identified mutations in *walK*, *yycH*, and *vraS*, which are consistent with our study, though *liaF* mutations were notably absent in their lines. This discrepancy could reflect differences in genetic background, as we evolved MSSA populations, whereas their study used MRSA. Thus, genetic background and environmental context together shape collateral responses, reinforcing the clinical relevance of considering both factors when interpreting experimental evolution studies.

Nafcillin and other antistaphylococcal penicillins are the current recommended treatments for MSSA bacteremia. Nonetheless, many clinicians support first-line cefazolin therapy instead because patients can better tolerate its side effects (39–42). Other clinicians prefer nafcillin because cefazolin is susceptible to cleavage by *S. aureus*-produced penicillinases and, therefore, may be less effective when there is a high bacterial titer (43, 44). We offer an evolutionary perspective on this debate: specific mutations that arise during empiric vancomycin therapy may negatively impact treatment effectiveness. On the one hand, *rpsU* mutations may predispose MSSA infections to suboptimal treatment with cefazolin or nafcillin, as even marginal resistance increases — such as those we observed experimentally — could become clinically significant if initial susceptibility is already close to clinical resistance thresholds. On the other hand, mutations in the WalKR regulon might promote favorable treatment outcomes. Although our results raise concerns about the reliability of these drugs as first-line agents in specific contexts, further research is needed to support these findings.

Our study, alongside corroborating evidence (25, 45), suggests that the evolution of vancomycin resistance may predictably impact collateral drug sensitivities, highlighting the key role that treatment history plays in shaping resistance outcomes. In this context, several studies have shown that appropriately designed sequential therapies hold promise for steering evolutionary trajectories to mitigate resistance (10, 26, 46, 47). Moreover, recent work that accounts for fluctuating environmental pressures (48) further illustrates how a nuanced understanding of evolution can inform treatment design. Together, these studies provide a conceptual framework that not only aligns with our study but also suggests promising strategies for integrating evolutionary insights into antimicrobial therapy.

### Using Collateral Response Scores to inform antibiotic treatment

When clinicians use antimicrobial susceptibility test results to guide treatment decisions, they participate in a decision-making process that involves uncertainty and risk. Results reflect the pathogen’s phenotypic state before empiric therapy and cannot account for adaptation during treatment, potentially leading to shifts in susceptibility. Our findings highlight this possibility. Confidence in the accuracy and relevance of initial susceptibility test results may, therefore, lead to ineffective treatment choices. It is paramount that clinicians have a well-informed, probabilistic understanding of how infections respond to treatment, informed by their prior evolutionary history.

We propose the Collateral Response Score (CRS) to address this issue. This composite metric accounts for the stochastic nature of evolution by quantifying the net collateral effect of antibiotic exposure. We defined it as,

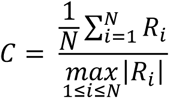

where *R*_*i*_ denotes the log_2_-fold change in MIC between replicate *ii* and the ancestor, *R*_*i*_ = log_2_(MIC_*ii*,evolved_/MIC_ancestor_), and *N* indicates the number of replicate populations.

The CRS is standardized and easily interpretable, yielding values between −1 and 1 that reflect the overall direction and magnitude of changes in antibiotic susceptibility relative to a prior genotypic state. Negative values indicate that, on average, evolved lines exhibit collateral sensitivity, and positive values indicate collateral resistance. A CRS close to −1 or 1 signifies substantial and probable changes in either direction, whereas values near zero denote only modest changes (or, importantly, balanced divergent changes).

This score is also amenable to statistical investigation. The tests discussed in the previous sections evaluated whether the distributions of MICs differed significantly between the ancestral clone and its derived VISA populations, or whether there was a correlation between specific genomic profiles and collateral response estimates. Nevertheless, these tests alone cannot provide information about the likelihood of a particular collateral response, nor its strength and direction. These attributes are essential information for clinicians making therapeutic decisions. Consider, for example, a situation where a susceptibility test indicates that an MSSA infection, previously treated with vancomycin, is sensitive to both cefazolin and nafcillin. How should a clinician tailor therapy, given that sensitivity might change with the evolution of *rpsU* mutations? In that spirit, we calculated bootstrap distributions of CRS, categorized by drug, pathway, and their interaction. The net collateral effect of vancomycin exposure on first-line drug susceptibility is significant if the bootstrap 95% confidence interval of a CRS estimate does not include zero.

Overall, the CRS distributions support our earlier findings while revealing important nuances in collateral responses. For example, although VISA populations exhibited the highest overall collateral sensitivity to gentamicin (**Fig. 3*C***), the considerable variability among replicates meant that individual populations could be nearly equally likely to evolve sensitivity or resistance (**Fig. 4*A***). The CRS effectively captures this variability, emphasizing its ability to represent both the magnitude and uncertainty inherent in evolutionary responses. Furthermore, mutations in *rpsU* modestly increased collateral resistance across drugs (**Fig. 4*B***), while mutations in the WalKR regulon and *rpsU* were associated with a higher likelihood of nafcillin collateral sensitivity and cross-resistance, respectively.

**Fig. 4.**
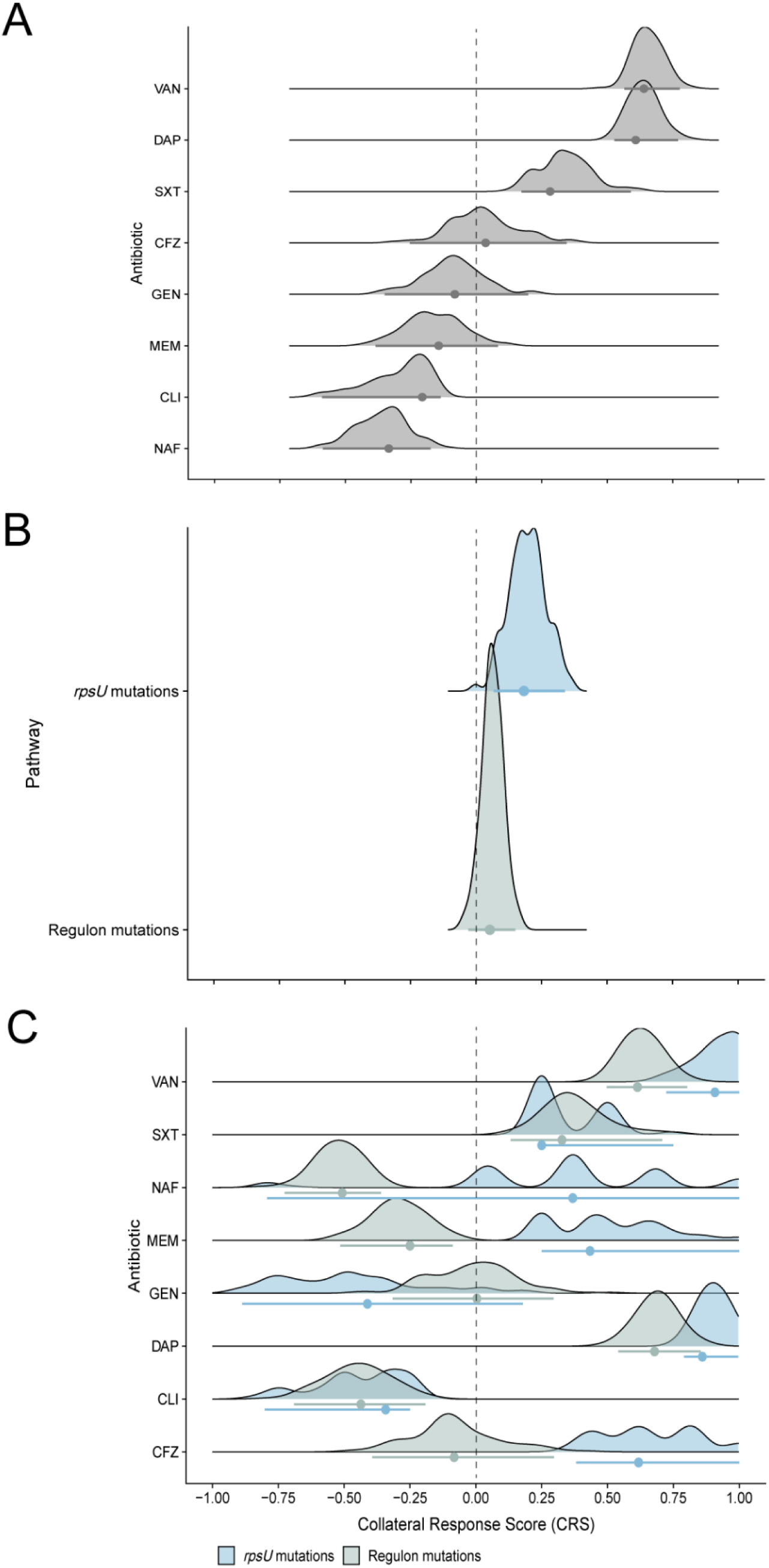
Bootstrap distributions of Collateral Response Scores (CRS) across **(*A*)** antibiotics, **(*B*)** genomic pathways, and their **(*C*)** interaction. The CRS is the mean log_2_-fold change in MIC between the evolved VISA populations and the ancestor, normalized by the maximum absolute change in MIC. Values range from –1 (uniform collateral sensitivity) to 1 (uniform collateral resistance). Each distribution represents 100 bootstrap replicates, with point estimates representing the mean CRS and error bars indicating the percentile-based 95% confidence interval. The dashed vertical line at zero denotes the threshold between collateral sensitivity and resistance.

Importantly, however, drug response outcomes might change in different experimental and clinical contexts. Here, we outline three possible future studies that address these contexts, generalize our findings, and possibly validate the CRS. First, one could examine whether *S. aureus* clinical isolates with different genetic backgrounds exhibit similar patterns of collateral sensitivity or resistance. A bacterium’s background may direct evolution toward some pathways while constraining others (49, 50), similar to how the evolution of WalKR regulon or *rpsU* mutations precluded the other in our MSSA lines. This process, known as historical contingency, may also influence how effectively a population compensates for increased susceptibility to first-line drugs through subsequent evolution (23, 51–53). Second, one might investigate how biofilm formation impacts collateral drug responses in MSSA and how different antibiotic delivery systems alter these patterns. These studies would bridge the gap between laboratory findings and clinical applicability, especially considering the relevance to implant-associated infections caused by biofilms and heart valve vegetations (54, 55). Third, we acknowledge that our experimental design, which involves progressively increasing vancomycin concentrations, differs from the dosing regimens typically used in clinical settings. This approach may have influenced the evolutionary trajectories observed in our study, potentially leading to the accumulation of mutations distinct from those that would occur under constant therapeutic doses. For example, Oz and colleagues showed that when *E. coli* lines evolve antibiotic resistance under strong selection, they exhibit more collateral resistance and have more mutations in drug pathway-specific genes than populations evolved under weak selection (56). While our method allowed us to explore a range of evolutionary responses, we acknowledge that our approach may limit the clinical relevance of our findings. Further studies under higher, clinically relevant doses (and over shorter durations) are warranted to understand the dynamics of resistance evolution in therapeutic contexts.

Nonetheless, our study reveals that vancomycin adaptation in MSSA proceeds through at least two distinct evolutionary pathways — one characterized by alterations in cell wall metabolism and the other by changes in global stress response. These adaptive trajectories result in contrasting collateral responses, with WalKR regulon mutations conferring enhanced sensitivity to nafcillin, while *rpsU* mutations tend to maintain or increase resistance to both nafcillin and cefazolin. Contingency highlights the importance of accounting for stochasticity when designing therapeutic strategies, underscoring that anticipating evolutionary variability may be crucial for preserving antibiotic efficacy and enhancing clinical outcomes.

## Materials and Methods

### Ancestral strain and experimental evolution

We used *Staphylococcus aureus* subsp. *aureus* Rosenbach (ATCC 29213) as the ancestor of our evolution experiment. This methicillin-susceptible strain was originally isolated from human pleural fluid in 1884 (57) and is an international quality control standard with defined susceptibilities to many antibiotics. According to the CLSI (58), vancomycin’s MIC on this strain is 0.5 – 2 µg/mL. Clinical *S. aureus* isolates with MICs <2 µg/mL are therefore considered susceptible to this drug. The clinical MIC breakpoints differentiating intermediate- and complete-resistant phenotypes are 4 – 8 and >16 µg/mL, respectively. We used these breakpoints to classify phenotypes in our study.

All experiments were performed at 37^°^C, unless otherwise noted. We revived the ATCC 29213 strain from a frozen stock by streaking cells onto tryptic soy agar (TSA) plates supplemented with 5% sheep blood (Remel, Lenexa, KS). We randomly chose isolated colonies from these plates to establish 18 replicate populations in tryptic soy broth (TSB) (Cleveland Clinic, Cleveland, OH). To start the evolution experiment, we prepared a linear series of vancomycin dilutions in TSB, ranging in concentrations from 0.25 to 1.5 µg/mL. Each replicate was aliquoted into equal volumes of the vancomycin-amended TSB in 96-well plates, resulting in a total dilution of 1:400, and incubated without shaking for 22 hours. Next, we transferred 1:400 cells from the highest concentration with visible growth for each replicate into fresh TSB with increased vancomycin concentrations. We transferred the replicate populations every 18–24 hours in the unshaken medium until they grew at concentrations between 4 and 8 µg/mL for two consecutive transfers. Replicates that did not grow at any concentration after 24 hours were left to incubate until they did so, but no longer than 96 hours. Additionally, we randomly chose 87 isolated colonies of the ATCC ancestor to establish replicate control populations. We propagated them for seven days under the same conditions, but in TSB without vancomycin. Undiluted cultures of the evolved populations were frozen at –80*^◦^*C in TSB supplemented with 15% glycerol as a cryoprotectant. The Cleveland Clinic Institutional Biosafety Committee approved all methods.

### Library preparation and whole-genome sequencing

Frozen glycerol stocks of ancestral and VISA population samples were cultured overnight in TSB with 1.5 µg/mL vancomycin, while the control populations were grown in vancomycin-free TSB. We centrifuged the overnight cultures at 8,000 rpm for 3 minutes and removed the supernatant from the bacterial pellets. SeqCenter (Pittsburgh, PA) prepared sample libraries from the ancestor and vancomycin-adapted lines using the Illumina DNA Prep kit and IDT 10bp UDI indices, followed by sequencing on an Illumina NextSeq 2000. Demultiplexing, quality control, and adapter trimming were performed using bcl-convert (v3.9.3).

SeqCoast Genomics (Portsmouth, NH) lysed the control samples using MagMAX Microbiome bead-beating tubes and extracted DNA using the Qiagen DNeasy 96 PowerSoil Pro QIAcube HT Kit. Samples were prepared for whole-genome sequencing using the Illumina DNA Prep tagmentation kit with Illumina Unique Dual Indexes. Sequencing was performed on the Illumina NextSeq 2000 platform using a 300-cycle flow cell kit to produce 2×150 bp paired-end reads. Moreover, 1 – 2% PhiX control was spiked into the run to support optimal base calling. Read demultiplexing and trimming, and run analytics were performed using DRAGEN v4.2.7, an onboard analysis software on the NextSeq 2000. All resulting FASTQ files of paired-end reads for the ancestor, control, and vancomycin-adapted populations were deposited in the NCBI Sequence Read Archive (accession no PRJNA1075422) (59).

### Mutation identification and statistical methodology

The sequencing reads were filtered to remove low-quality bases using Trimmomatic v0.39 (60). We clipped the reads when the average quality score was <20 in a 4-bp window and to a minimum length of 36 bp. Next, we used *breseq* v0.39.0 (61) to identify mutations in two steps. First, we used this bioinformatic pipeline with default parameters to map the ancestral strain reads to the ATCC 29213 reference genome. This step accounted for mutations present at the beginning of our evolution experiment. Second, we applied these background mutations to the ATCC 29213 genome and reran *breseq* in polymorphism mode to map the evolved population reads to this updated reference.

We manually curated the sequencing results by excluding mutations with frequencies below 5% and within multicopy elements, such as ribosomal RNA operons and synthetases responsible for charging tRNA. Gene conversions may cause these mutations, but short-read sequencing data cannot fully resolve them (24). We chose the 5% frequency threshold given the high genomic coverage of our samples. Moreover, we included only those “qualifying” mutations that unambiguously impact a single gene (24, 62). These include nonsynonymous point mutations, small indels that occur within genes, mutations within 150 bp upstream of the start of a gene, and large deletions if at least one affected gene was mutated in another population.

We calculated the similarity of gene-level mutations between independent lines to evaluate genomic parallelism in the VISA populations relative to control lines. For each population pair, we calculated Dice’s similarity coefficient, *S*, as,

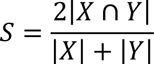

where |*X*| and |*Y*| represent the sets of genes with qualifying mutations in each population and |*X* ∩ *Y*| denotes the number of genes mutated in both populations. A value of zero indicates that the pair shares no mutations in common, whereas a value of one indicates complete overlap (24, 62, 63). We calculated *S* for every pair of populations within the vancomycin and control conditions, and then derived the average pairwise similarity for each group. The difference between the average similarity in the vancomycin-adapted lines (*S*_*v*_) and in the control lines (*S*_*c*_) served as the test statistic; a positive difference indicates that the VISA lines exhibit greater genomic parallelism than the control lines.

To determine the statistical significance of this difference, we employed a permutation test. In each of the 10,000 permutations, the populations were randomly reassigned to the vancomycin or control groups, and the difference *S*_*v*_ − *S*_*c*_ was recalculated, thereby generating an empirical distribution under the null hypothesis of equal genomic parallelism. We then obtained an approximate *P*-value from the proportion of permutations with a test statistic at least as extreme as the observed value.

Next, we used penalized logistic regression with L1 (LASSO) and L2 (Ridge) regularization to identify genes associated with vancomycin selection. Given the high dimensionality of the initial mutation set, we performed feature selection in two steps. First, we assessed the convergence of model coefficient estimates over 1,000 iterations. In each iteration, the optimal regularization strength (λ) and mixing parameters (α) were determined via three-fold cross-validation to minimize deviance, and a multivariate logistic regression model was fitted using these parameters. We only included in our model qualifying mutations that occurred in at least three populations, resulting in 446 gene-level mutations. For every qualifying mutation, we computed the cumulative mean of its coefficient to assess whether the point estimate stabilized as the number of iterations increased. In parallel, we calculated the cumulative standard error, reflecting the estimate’s precision across iterations. We considered convergence achieved at iteration 250, as the cumulative mean (**Fig. S3*A***) and standard error (**Fig. S3*B***) exhibited a marginal change with further iterations. Second, we performed the penalized logistic regression procedure as before with 250 iterations. Given the sparsity-inducing property of L1 regularization, we tracked the frequency of non-zero coefficients across iterations. Mutations exhibiting non-zero coefficients in at least 20% of the iterations were retained, constituting the final set of informative features for subsequent Bayesian latent class analysis (BLCA) (SI Methods). We computed the 95% confidence intervals for each gene’s coefficient and then exponentiated the endpoints to obtain the confidence intervals for the odds ratio. We provide the datasets and details of our statistical analyses in an R Notebook on GitHub (64).

### Estimating collateral drug responses

We estimated the collateral responses of the 18 evolved VISA populations to first-line antibiotics using the broth microdilution method outlined by the CLSI (58). We supplemented Mueller-Hinton broth (MHB) (Fisher Bioreagents, Ottawa, CA) with 20–25 mg/L Ca^2+^ (CaCl_2_ • 2H_2_O, Fisher Bioreagents, Ottawa, CA) and 10–15 mg/L Mg^2+^ (MgCl_2_, Invitrogen, Vilnius, LT). We used this cation-adjusted MHB (CA-MHB) to test the susceptibilities of the evolved populations to cefazolin, clindamycin, daptomycin, gentamycin, meropenem, nafcillin, and trimethoprim/sulfamethoxazole. For daptomycin and nafcillin susceptibility tests, we used CA-MHB that was additionally supplemented with 25 mg/L Ca^2+^ (for a total calcium concentration of 50 mg/L) and 2% (w/v) NaCl (Fisher Bioreagents, Ottawa, CA), respectively (65). CA-MHB was stored at 4*^◦^*C to prevent ion precipitation.

We inoculated 100 µL of cells from frozen samples into CA-MHB with 1 µg/mL vancomycin (i.e., the MIC of the ancestral strain) to maintain resistance. After overnight growth, the cultures were diluted to a McFarland 0.5 standard and then again by 200-fold in MHB. We aliquoted an equal volume of these cells across a series of linear concentrations of a given antibiotic in MHB. These concentrations ranged from 0.125 to 3× the median MIC of the wild-type *S. aureus* ATCC 29213 clone. To meet the CLSI standard (58), we incubated the vancomycin and nafcillin cultures for 24 hours at 35*^◦^*C. The other cultures were incubated at 37*^◦^*C for 18–20 hours. The MIC of each sample was the lowest antibiotic concentration that inhibited visual growth.

We quantified the susceptibility of three technical replicates for each of the 18 evolved VISA populations across the seven antibiotics, totaling 378 MIC measurements. In addition, we included the ancestor in each broth microdilution plate as a control and reference. The susceptibility of this strain was estimated using eight technical replicates per drug, resulting in 56 MIC measurements. We calculated the collateral drug response, *R*, for each technical replicate, *i*, from population, *j*, as:

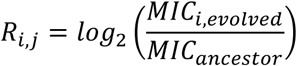

We analyzed the average collateral response values among the three technical replicates for each population.

## Acknowledgments

We thank the members of Theory Division for their valuable discussions and the Cleveland Clinic Microbiology Department for providing the ATCC 29213 strain. We acknowledge financial support from an HHMI Hanna H. Gray Fellowship (to K.J.C.), a Ruth L. Kirschstein NRSA Institutional Research Training Grant (to J.M.), and an NIH R37 grant (5R37CA244613-02) (to J.G.S.).

## Author Contributions

K.J.C., D.C., and J.G.S. conceived of and designed the study. K.J.C., D.C., J.C., and A.S. performed the experiments. K.J.C., D.C., A.D., and J.G. analyzed all data. K.J.C. and A.D. wrote the associated analysis script and prepared figures. K.J.C. and J.G.S. supervised the project. K.J.C., D.C., and A.D. wrote the original draft. K.J.C., D.C., A.D., R.A.B., Z.D.C.B., and J.G.S. reviewed and edited the paper. All authors approved the final version.

## Competing Interests

The authors declare no competing interests.

## Data Availability

We provide the datasets and analysis code on GitHub (https://github.com/KyleCard/S_aureus_evolution) (64). Sequence read data have been deposited in the National Center for Biotechnology Information Sequence Read Archive (accession no PRJNA1075422) (59).

## SI Methods

Following the feature selection by multivariate logistic regression, we performed Bayesian latent class analysis (BLCA) using the R package BayesLCA (1) with selected features (genes) mutated in at least three lines to identify distinct evolutionary trajectories. In this framework, the posterior distribution of parameters given the observations can be written as,

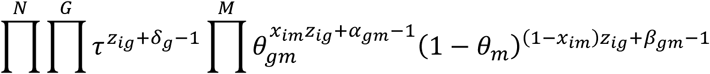

where τ is the estimate for the population frequency of a particular group, θ is a GxM dimensional matrix of mutation probability given the group assignment, and δ, α, and β are parameters for the conjugate prior distributions for the Dirichlet and Beta distributions, respectively. We set the δ parameter to 1.0 as a uniform prior over class distributions and α and β to 0.5 to increase density around 0.0 and 1.0. Given the computational constraints of Markov chain Monte Carlo-based estimation, we used the expectation-maximization algorithm. To further improve the robustness of the clustering, we used the BLCA over 1000 iterations, where a random 90% subsampling was performed for both features and observations at each iteration. The subsampled data was clustered with *k* set from 1 to 5, and the best number of clusters was selected using AIC. At each clustering iteration, we tracked which observations were clustered together. Subsequently, we created a consensus matrix quantifying the frequency of pairwise observations, and a final clustering was performed using hierarchical clustering with Ward’s criteria on the frequency matrix.

## SI Figures

**Fig. S1.**
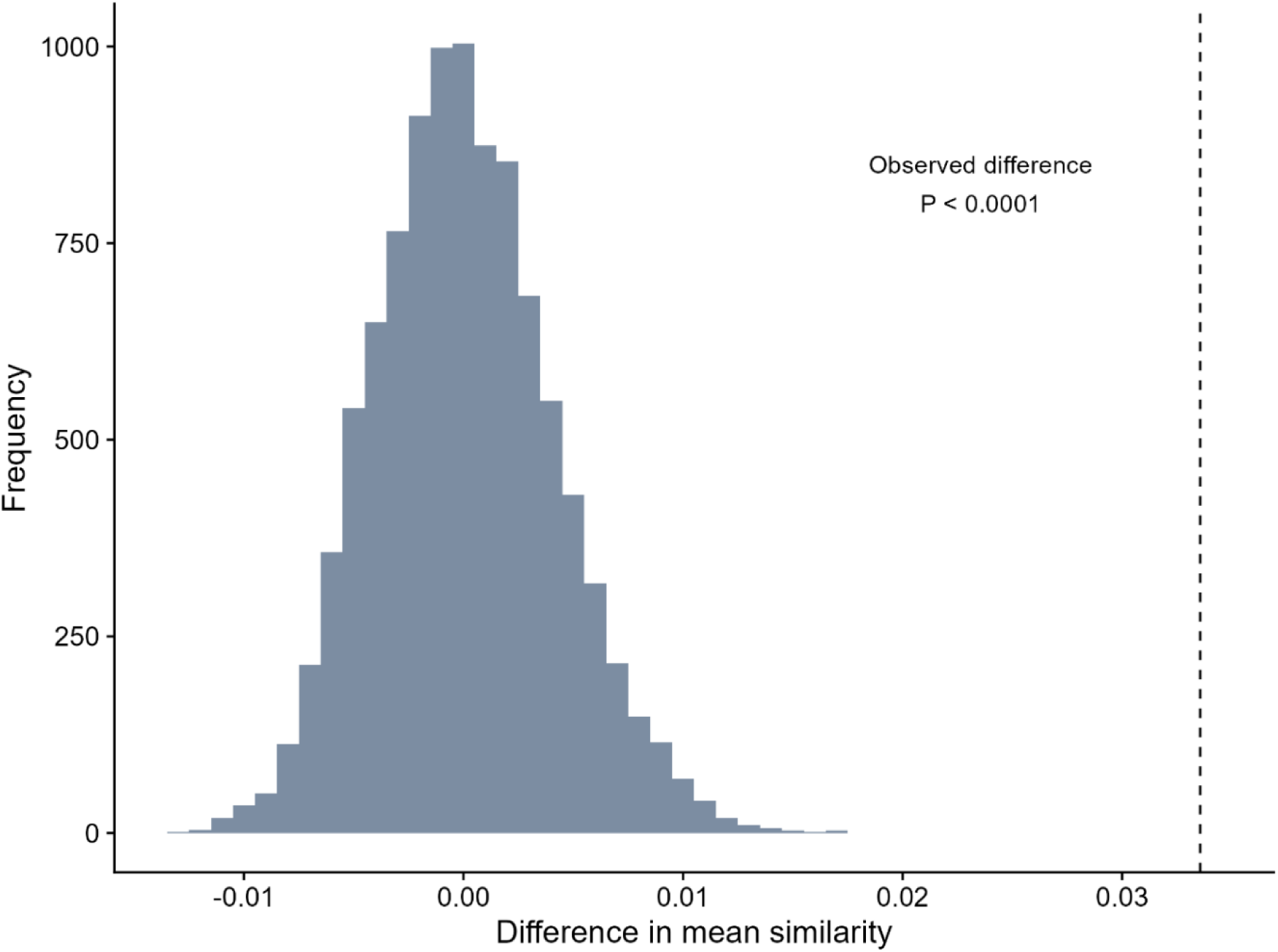
The null distribution of the difference in mean gene-level similarity between vancomycin-adapted and control populations. We calculated Dice’s similarity coefficients for each population pair using only the qualifying mutations that unambiguously affected single genes. We assessed the difference in average similarity between the vancomycin-adapted and control lines. Then, we shuffled the population labels randomly between the two treatment groups and recalculated the difference in mean similarity. This process was repeated 10,000 times. The resulting distribution of differences is presented here. The observed difference of 0.034 surpassed all permutations (*P* < 0.0001), indicating a significantly higher level of genomic parallelism under vancomycin selection.

**Fig. S2.**
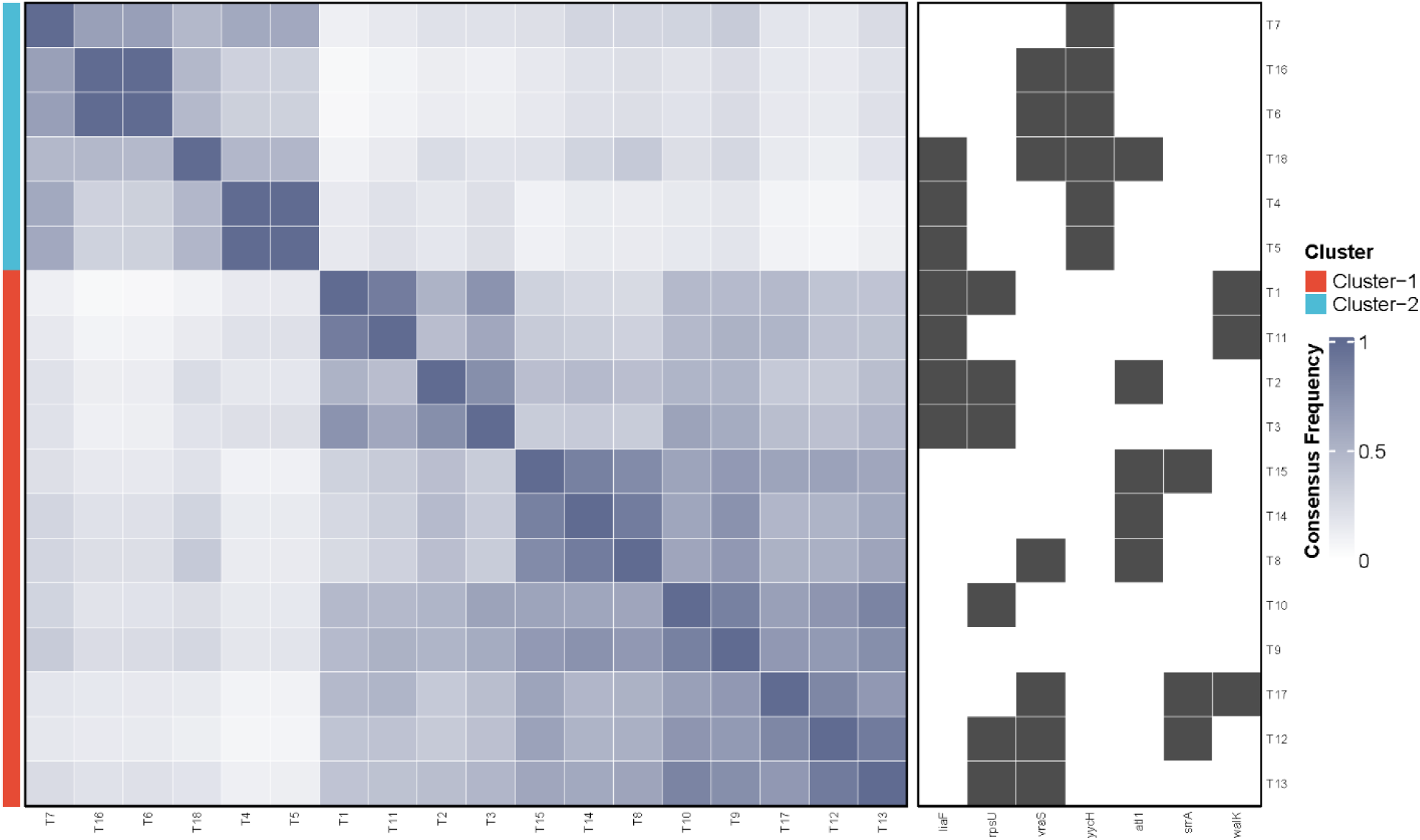
Two distinct adaptive trajectories occurred under vancomycin selection. The heatmap represents the consensus matrix, generated through iterative Bayesian latent class analysis (BLCA) with random subsampling of observations and genomic features (Supplemental Methods), indicating the presence of two distinct clusters that correspond to unique genomic profiles.

**Fig. S3.**
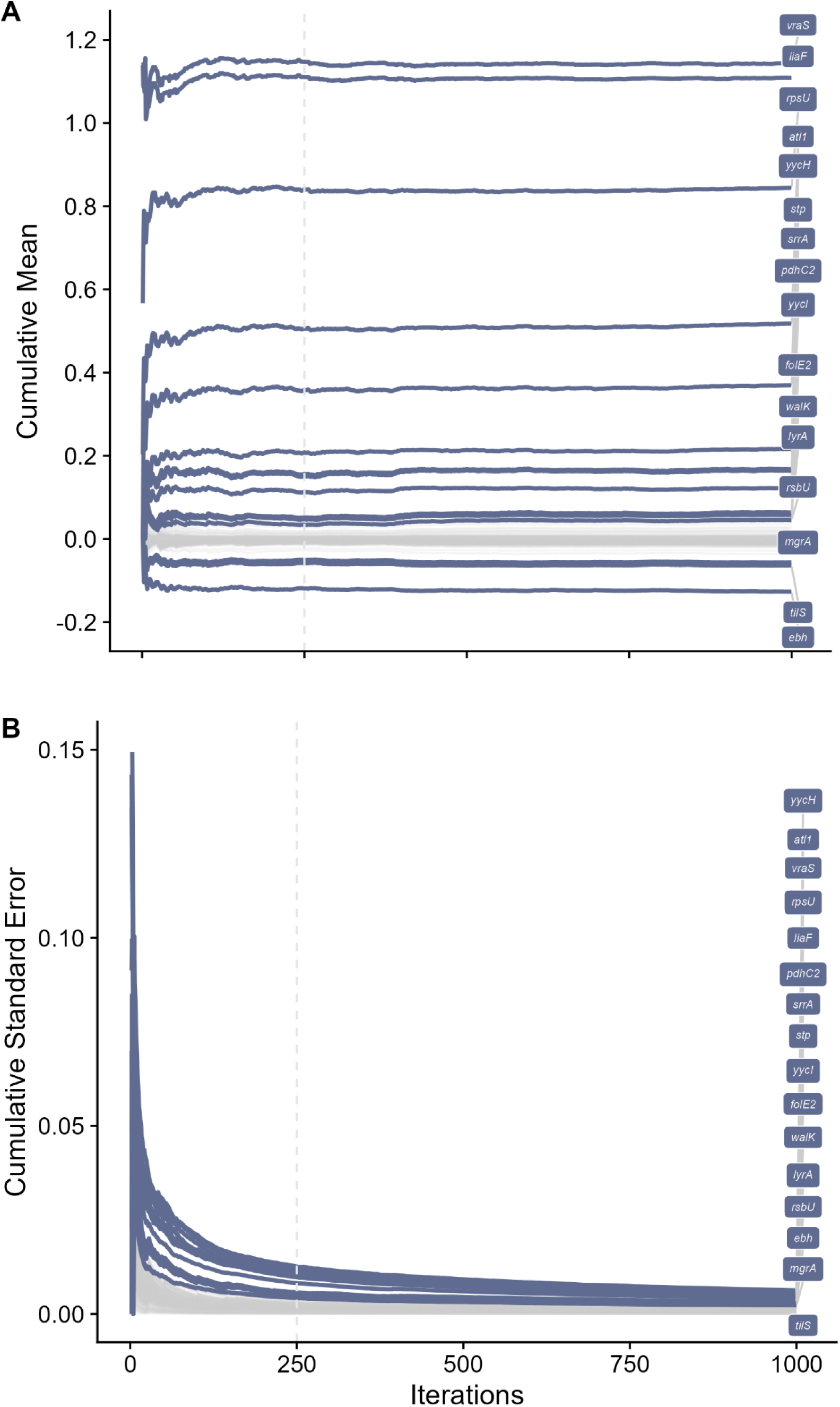
Tests for convergence in logistic regression model coefficients. Over 1,000 iterations of a penalized regression procedure (Materials and Methods), we estimated (***A***) the cumulative mean and (***B***) the standard error for coefficients corresponding to qualifying gene-level mutations. Blue highlighted lines denote genes with non-zero coefficients in at least 20% of the iterations. Labels are ordered by decreasing cumulative mean and standard error. Grey highlighted lines indicate all other genes. The dotted line indicates the iteration where convergence is achieved.

## Notes

https://github.com/KyleCard/S_aureus_evolution

### Competing Interest Statement

The authors have declared no competing interest.

### Summary of Updates

We have included an additional experiment. In brief, we established 87 replicate populations from the S. aureus ATCC 29213 ancestor and transferred these populations for 7 days in vancomycin-free tryptic soy broth (TSB). We then sequenced these populations and compared these data with those from the experimental lines. We performed four additional analyses: (1) gene-level similarity with an associated permutation test, (2) penalized multivariate logistic regression with vancomycin treatment as the outcome and 446 gene-level mutations as predictors, (3) Bayesian latent-class analysis (BLCA) of the vancomycin-treated lines, and (4) correlation analysis (i.e., point biserial tests). Lastly, we developed a new metric, the Collateral Response Score, to quantify the probability and magnitude of collateral responses. As such, we have: 1. Added a new Figure 2, which reports the mutated genes and pathways in the experimentally evolved VISA populations. This figure includes the 16 genes with non-zero odds ratios, as determined from the regression model (i.e., these genes are associated with either vancomycin or permissive conditions), along with a heatmap of these gene-level mutations. 2. Added a new Figure 3 reporting collateral responses stratified by pathway targeted under vancomycin selection. Mutations in the WalKR regulon were associated with increased nafcillin collateral sensitivity, whereas mutations in rpsU were associated with cross-resistance. 3. Added Figure 4, which shows the bootstrapped distributions of Collateral Response Scores (CRS) across antibiotics, between pathways, and their interaction. We extended the concept of the likelihood estimate in the original draft to derive the CRS. Briefly, it is a standardized metric that yields values between -1 and 1, reflecting the overall direction and magnitude of changes in antibiotic susceptibility relative to a prior genotypic state. 4. Added Figure S1, which reports the distribution of permuted differences in the mean similarity between vancomycin-adapted and control groups, and indicates where the observed test statistic falls on this distribution. 5. Added Figure S2, which reports the BLCA results. 6. Added Figure S3 on logistic model convergence estimates.

## References

1. T. L. Holland, C. Arnold, V. G. Fowler Jr, Clinical management of *Staphylococcus aureus* bacteremia: A review. JAMA 312, 1330–1341 (2014).

2. L. M. Baddour, et al., Infective endocarditis in adults: Diagnosis, antimicrobial therapy, and management of complications: A scientific statement for healthcare professionals from the American Heart Association. Circulation 132, 1435–1486 (2015).

3. S. Y. C. Tong, J. S. Davis, E. Eichenberger, T. L. Holland, V. G. Fowler Jr, *Staphylococcus aureus* infections: Epidemiology, pathophysiology, clinical manifestations, and management. Clin. Microbiol. Rev. 28, 603–661 (2015).

4. T. J. Cahill, et al., Antibiotic prophylaxis for infective endocarditis: A systematic review and meta-analysis. Heart 103, 937–944 (2017).

5. B. P. Howden, et al., Evolution of multidrug resistance during *Staphylococcus aureus* infection involves mutation of the essential two component regulator WalKR. PLoS Pathog. 7, e1002359 (2011).

6. S. Gardete, A. Tomasz, Mechanisms of vancomycin resistance in *Staphylococcus aureus*. J. Clin. Invest. 124, 2836–2840 (2014).

7. Q. Hu, H. Peng, X. Rao, Molecular events for promotion of vancomycin resistance in vancomycin intermediate *Staphylococcus aureus*. Front. Microbiol. 7, 1601 (2016).

8. Y. Cong, S. Yang, X. Rao, Vancomycin resistant *Staphylococcus aureus* infections: A review of case updating and clinical features. J. Adv. Res. 21, 169–176 (2020).

9. C. Cervera, et al., Effect of vancomycin minimal inhibitory concentration on the outcome of methicillin-susceptible *Staphylococcus aureus* endocarditis. Clin. Infect. Dis. 58, 1668–1675 (2014).

10. J. Maltas, K. B. Wood, Pervasive and diverse collateral sensitivity profiles inform optimal strategies to limit antibiotic resistance. PLoS Biol. 17, e3000515 (2019).

11. D. Nichol, et al., Antibiotic collateral sensitivity is contingent on the repeatability of evolution. Nat. Commun. 10, 334 (2019).

12. S. Heidarian, A. Guliaev, H. Nicoloff, K. Hjort, D. I. Andersson, High prevalence of heteroresistance in *Staphylococcus aureus* is caused by a multitude of mutations in core genes. PLoS Biol. 22, e3002457 (2024).

13. P. Suntharalingam, M. D. Senadheera, R. W. Mair, C. M. Lévesque, D. G. Cvitkovitch, The LiaFSR system regulates the cell envelope stress response in *Streptococcus mutans*. J. Bacteriol. 191, 2973–2984 (2009).

14. E. Galbusera, et al., Site-specific mutation of *Staphylococcus aureus* VraS reveals a crucial role for the VraR-VraS sensor in the emergence of glycopeptide resistance. Antimicrob. Agents Chemother. 55, 1008–1020 (2011).

15. C. Hafer, Y. Lin, J. Kornblum, F. D. Lowy, A.-C. Uhlemann, Contribution of selected gene mutations to resistance in clinical isolates of vancomycin-intermediate *Staphylococcus aureus*. Antimicrob. Agents Chemother. 56, 5845–5851 (2012).

16. J. Koomen, et al., Ribosomal mutations enable a switch between high fitness and high stress resistance in *Listeria monocytogenes*. Front. Microbiol. 15 (2024).

17. M. D. S. Basco, et al., Reduced vancomycin susceptibility and increased macrophage survival in *Staphylococcus aureus* strains sequentially isolated from a bacteraemic patient during a short course of antibiotic therapy. J. Med. Microbiol. 68, 848–859 (2019).

18. S. Dubrac, I. G. Boneca, O. Poupel, T. Msadek, New insights into the WalK/WalR (YycG/YycF) essential signal transduction pathway reveal a major role in controlling cell wall metabolism and biofilm formation in *Staphylococcus aureus*. J. Bacteriol. 189, 8257–8269 (2007).

19. M. Gajdiss, et al., YycH and YycI regulate expression of *Staphylococcus aureus* autolysins by activation of WalRK phosphorylation. Microorganisms 8, 870 (2020).

20. J. E. Sulaiman, L. Wu, H. Lam, Mutation in the two-component system regulator YycH leads to daptomycin tolerance in methicillin-resistant *Staphylococcus aureus* upon evolution with a population bottleneck. *Microbiol*. Spectrum 10, e01687–22 (2022).

21. S. Dubrac, T. Msadek, Identification of genes controlled by the essential YycG/YycF two-component system of *Staphylococcus aureus*. J. Bacteriol. 186, 1175–1181 (2004).

22. A. Delauné, et al., The WalKR system controls major staphylococcal virulence genes and is involved in triggering the host inflammatory response. Infect. Immun. 80, 3438–3453 (2012).

23. M. Lukačišinová, B. Fernando, T. Bollenbach, Highly parallel lab evolution reveals that epistasis can curb the evolution of antibiotic resistance. Nat. Commun. 11, 3105 (2020).

24. K. J. Card, M. D. Thomas, J. L. Graves, J. E. Barrick, R. E. Lenski, Genomic evolution of antibiotic resistance is contingent on genetic background following a long-term experiment with *Escherichia coli*. Proc. Natl. Acad. Sci. U.S.A. 118, e2016886118 (2021).

25. A. Fait, et al., Adaptive laboratory evolution and independent component analysis disentangle complex vancomycin adaptation trajectories. Proc. Natl. Acad. Sci. U.S.A. 119, e2118262119 (2022).

26. D. T. Weaver, E. S. King, J. Maltas, J. G. Scott, Reinforcement learning informs optimal treatment strategies to limit antibiotic resistance. Proc. Natl. Acad. Sci. U.S.A. 121, e2303165121 (2024).

27. G. R. Corey, *Staphylococcus aureus* bloodstream infections: Definitions and treatment. Clin. Infect. Dis. 48, S254–S259 (2009).

28. V. Delgado, et al., 2023 ESC guidelines for the management of endocarditis. Eur. Heart J. 44, 3948–4042 (2023).

29. Y. Benjamini, Y. Hochberg, Controlling the false discovery rate: A practical and powerful approach to multiple testing. J. R. Stat. Soc. B. 57, 289–300 (1995).

30. CLSI supplement M100, M100: Performance standards for antimicrobial susceptibility testing, 35th Ed. (Clinical Laboratory Standards Institute, 2025).

31. L. Cui, E. Tominaga, H. Neoh, K. Hiramatsu, Correlation between reduced daptomycin susceptibility and vancomycin resistance in vancomycin-intermediate *Staphylococcus aureus*. Antimicrob. Agents Chemother. 50, 1079–1082 (2006).

32. P. G. Kelley, W. Gao, P. B. Ward, B. P. Howden, Daptomycin non-susceptibility in vancomycin-intermediate *Staphylococcus aureus* (VISA) and heterogeneous-VISA (hVISA): Implications for therapy after vancomycin treatment failure. J. Antimicrob. Chemother. 66, 1057–1060 (2011).

33. C. A. Arias, et al., Genetic basis for in vivo daptomycin resistance in enterococci. N. Engl. J. Med. 365, 892–900 (2011).

34. J. M. Munita, et al., Correlation between mutations in *liaFSR* of *Enterococcus faecium* and MIC of daptomycin: Revisiting daptomycin breakpoints. Antimicrob. Agents Chemother. 56, 4354–4359 (2012).

35. G. Sakoulas, J. Alder, C. Thauvin-Eliopoulos, R. C. Moellering, G. M. Eliopoulos, Induction of daptomycin heterogeneous susceptibility in *Staphylococcus aureus* by exposure to vancomycin. Antimicrob. Agents Chemother. 50, 1581–1585 (2006).

36. C. Watanakunakorn, Clindamycin therapy of *Staphylococcus aureus* endocarditis: Clinical relapse and development of resistance to clindamycin, lincomycin and erythromycin. Am. J. Med. 60, 419–425 (1976).

37. H. Tissot-Dupont, et al., High-dose trimethoprim-sulfamethoxazole and clindamycin for *Staphylococcus aureus* endocarditis. Int. J. Antimicrob. Agents 54, 143–148 (2019).

38. H. Machado, et al., Environmental conditions dictate differential evolution of vancomycin resistance in *Staphylococcus aureus*. *Commun*. Biol. 4, 793 (2021).

39. M. L. Schweizer, et al., Comparative effectiveness of nafcillin or cefazolin versus vancomycin in methicillin-susceptible *Staphylococcus aureus* bacteremia. BMC Infect. Dis. 11, 279 (2011).

40. J. Li, et al., Comparison of cefazolin versus oxacillin for treatment of complicated bacteremia caused by methicillin-susceptible *Staphylococcus aureus*. Antimicrob. Agents Chemother. 58, 5117–5124 (2014).

41. I. Youngster, E. S. Shenoy, D. C. Hooper, S. B. Nelson, Comparative evaluation of the tolerability of cefazolin and nafcillin for treatment of methicillin-susceptible *Staphylococcus aureus* infections in the outpatient setting. Clin. Infect. Dis. 59, 369–375 (2014).

42. A. M. Gandhi, M. D. Shah, L. E. Donohue, H. L. Cox, J. C. Eby, Tolerability of cefazolin in nafcillin-intolerant patients for the treatment of methicillin-susceptible *Staphylococcus aureus* infections. Clin. Infect. Dis. 73, 1650–1655 (2021).

43. E. C. Nannini, et al., Determination of an inoculum effect with various cephalosporins among clinical isolates of methicillin-susceptible *Staphylococcus aureus*. Antimicrob. Agents Chemother. 54, 2206–2208 (2010).

44. K. V. Singh, et al., Efficacy of ceftaroline against methicillin-susceptible *Staphylococcus aureus* exhibiting the cefazolin high-inoculum effect in a rat model of endocarditis. Antimicrob. Agents Chemother. 61, 10.1128/aac.00324-17 (2017).

45. M. Su, et al., Effect of genetic background on the evolution of vancomycin-intermediate *Staphylococcus aureus* (VISA). PeerJ 9, e11764 (2021).

46. D. Nichol, et al., Steering evolution with sequential therapy to prevent the emergence of bacterial antibiotic resistance. PLoS Comput. Biol. 11, e1004493 (2015).

47. S. Iram, et al., Controlling the speed and trajectory of evolution with counterdiabatic driving. Nat. Phys. 17, 135–142 (2021).

48. E. S. King, et al., Fitness seascapes are necessary for realistic modeling of the evolutionary response to drug therapy. bioRxiv 2022.06.10.495696 (2025). 10.1101/2022.06.10.495696.

49. Z. D. Blount, C. Z. Borland, R. E. Lenski, Historical contingency and the evolution of a key innovation in an experimental population of *Escherichia coli*. Proc. Natl. Acad. Sci. U.S.A. 105, 7899–7906 (2008).

50. Z. D. Blount, R. E. Lenski, J. B. Losos, Contingency and determinism in evolution: Replaying life’s tape. Science 362, eaam5979 (2018).

51. K. J. Card, T. LaBar, J. B. Gomez, R. E. Lenski, Historical contingency in the evolution of antibiotic resistance after decades of relaxed selection. PLoS Biol. 17, e3000397 (2019).

52. A. Santos-Lopez, et al., The roles of history, chance, and natural selection in the evolution of antibiotic resistance. eLife 10, e70676 (2021).

53. A. Fait, D. I. Andersson, H. Ingmer, Evolutionary history of *Staphylococcus aureus* influences antibiotic resistance evolution. Curr. Biol. 33, 3389–3397.e5 (2023).

54. P. S. Stewart, J. W. Costerton, Antibiotic resistance of bacteria in biofilms. Lancet 358, 135– 138 (2001).

55. K. Schilcher, A. R. Horswill, Staphylococcal biofilm development: Structure, regulation, and treatment strategies. Microbiol. Mol. Biol. Rev. 84, 10.1128/mmbr.00026-19 (2020).

56. T. Oz, et al., Strength of selection pressure is an important parameter contributing to the complexity of antibiotic resistance evolution. Mol. Biol. Evol. 31, 2387–2401 (2014).

57. A. Shiroma, et al., First complete genome sequences of *Staphylococcus aureus* subsp. *aureus* Rosenbach 1884 (DSM 20231T), determined by PacBio single-molecule real-time technology. Genome Announc. 3, 10.1128/genomea.00800-15 (2015).

58. CLSI, M07: Methods for dilution antimicrobial susceptibility tests for bacteria that grow aerobically, 12th Ed. (Clinical Laboratory Standards Institute, 2024).

59. K. J. Card, et al., Data from “Evolution under vancomycin selection drives divergent collateral sensitivity patterns in *Staphylococcus aureus*.” NCBI Sequence Read Archive. https://www.ncbi.nlm.nih.gov/bioproject/PRJNA1075422. Deposited 31 January 2025.

60. A. M. Bolger, M. Lohse, B. Usadel, Trimmomatic: a flexible trimmer for Illumina sequence data. Bioinformatics 30, 2114–2120 (2014).

61. J. E. Barrick, et al., Identifying structural variation in haploid microbial genomes from short-read resequencing data using *breseq*. BMC Genomics 15, 1039 (2014).

62. D. E. Deatherage, J. L. Kepner, A. F. Bennett, R. E. Lenski, J. E. Barrick, Specificity of genome evolution in experimental populations of *Escherichia coli* evolved at different temperatures. Proc. Natl. Acad. Sci. U.S.A. 114 (2017).

63. R. R. Sokal, F. J. Rohlf, Biometry: The Principles and Practices of Statistics in Biological Research, 3rd Ed. (W. H. Freeman and Company, 1994).

64. K. J. Card, et al., Data from “Evolution under vancomycin selection drives divergent collateral sensitivity patterns in *Staphylococcus aureus*.” GitHub. https://github.com/KyleCard/S_aureus_evolution. Deposited 2 April 2025.

65. I. Wiegand, K. Hilpert, R. E. W. Hancock, Agar and broth dilution methods to determine the minimal inhibitory concentration (MIC) of antimicrobial substances. Nat. Protoc. 3, 163–175 (2008).

## SI Reference

1. A. White, T. B. Murphy, BayesLCA: An R package for Bayesian latent class analysis. J. Stat. Softw. 16, 1–28 (2014).

